# Structure-based prediction and characterization of photo-crosslinking in native protein-RNA complexes

**DOI:** 10.1101/2022.06.02.494568

**Authors:** Huijuan Feng, Xiang-Jun Lu, Linxi Liu, Dmytro Ustianenko, Chaolin Zhang

**Affiliations:** Department of Systems Biology, Department of Biochemistry and Molecular Biophysics, Center for Motor Neuron Biology and Disease, Columbia University, New York NY 10032, USA; Department of Biological Sciences, Columbia University, New York, NY 10027, USA; Department of Statistics, Columbia University, New York NY 10027, USA

## Abstract

UV-crosslinking of protein and RNA in direct contacts has been widely used to study protein-RNA complexes despite our poor understanding on the mechanisms of photo-crosslinking. This knowledge gap is due to the challenge to precisely map the crosslink sites in protein and RNA simultaneously in their native sequence and structural contexts. Here we developed PxR3D-map, a computational method to analyze protein-RNA interactions and photo-crosslinking by bridging crosslinked nucleotides and amino acids mapped using different assays with protein-RNA complex structures. PxR3D-map reliably predicts crosslink sites using structural information characterizing protein-RNA interaction interfaces. We found that photo-crosslinking is facilitated by base stacking with not only aromatic residues, but also dipeptide bonds that involve glycine, and distinct mechanisms are utilized by different types of RNA-binding domains. Our work suggests protein-RNA photocrosslinking is highly selective in the cellular environment, which can guide interpretation of data generated by UV-crosslinking-based assays and further technology development.

## Introduction

Control of RNA metabolism, which is a critical component of gene expression regulation, relies on specific sequence or structure elements embedded in transcripts that are recognized by RNA-binding proteins (RBPs)^1, 2^. Such regulatory elements are frequently short (3-7 nucleotides) and degenerate, and protein-RNA interactions are highly dynamic, so that it remains challenging to understand how RBPs specifically recognize their targets^3–5^. A widely used approach to investigate protein-RNA interactions is to crosslink protein and RNA in direct contacts using UV, which induces covalently linked conjugates between the interacting amino acids and nucleotides^6–8^. UV crosslinking can be performed using tissues or cultured cells and crosslinked protein-RNA complexes can then be analyzed to identify both the RNA and protein components by different strategies. For example, crosslinking and immunoprecipitation (CLIP) is now the *de facto* standard method to isolate RNA fragments crosslinked to a particular RBP of interest followed by deep sequencing to map RBP binding footprints on a genome-wide scale^9–11^. Alternatively, RBPs can be pulled down through crosslinked RNA to recover their identities using a method named RNA-interactome capture^12–14^. These efforts have greatly expanded the list of RBPs and our understanding of how they contribute to RNA regulation.

Despite the wide applications of UV crosslinking-based assays, the biophysical basis of protein-RNA crosslinking is currently poorly understood, especially for native complexes *in vivo* or in cellular contexts. Earlier studies using *in vitro* crosslinking model systems that involve single amino acids (or dipeptides) with homopolynucleotide chains suggest that each of the RNA bases is capable of forming conjugates with a variety of amino acids or peptides^15, 16^, but these studies provide limited information about photo-crosslinking of macromolecular complexes in cells. To improve the resolution of protein-RNA interaction mapping, approaches have been developed to pinpoint the exact crosslinked amino acid or nucleotide at single-residue resolution, taking advantage of the covalently linked amino acid-nucleotide adducts. Previously, we developed computational methods to map the exact crosslink sites in RNA through analysis of crosslink-induced mutation sites (CIMS) or truncation sites (CITS) using CLIP data^17–19^. CIMS and CITS provide signatures of protein-RNA crosslinking introduced by interference of reverse transcription by the amino acid-RNA adducts. The RNA-interactome capture workflow has also been refined to map the crosslinked amino acids by considering the mass shift caused by the RNA moieties conjugated to the crosslinked peptides that are subject to massspectrometry analysis^20–22^. However, to our knowledge no current technologies can map crosslinked amino acid and nucleotide simultaneously in native protein-RNA complexes, although such efforts have been made for reconstituted protein-RNA complexes *in vitro* in a few cases^23–25^. As a consequence, it remains unclear how selective photo-crosslinking is or whether particular types of amino acid-nucleotide contacts are required for photo-adducts formation. Nevertheless, such knowledge is highly relevant for understanding protein-RNA complex structures and interpreting data generated by UV crosslinking-based assays.

We reasoned that the sequence and structural contexts of protein-RNA crosslink sites can be deduced by integrating the crosslinked nucleotides in RNA inferred by CIMS and CITS analysis and crosslinked amino acids identified in RNA-interactome capture with experimentally determined protein-RNA complex structures. In this work, we developed a computational method named PxR3D-map for this purpose. PxR3D-map systematically analyzes various structural features associated with each nucleotide and amino acid in direct contact in 3D protein-RNA complex structures, which are then used to classify crosslinked vs. non-crosslinked nucleotides, as well as crosslinked vs. non-crosslinked amino acids. More importantly, PxR3D-map ranks structural features based on their importance for classification, which provides mechanistic insights into the biophysical basis of protein-RNA photo-crosslinking and adduct formation in their native complexes.

## Results

### PxR3D-map method overview

The central idea of PxR3D-map is to overlay UV crosslink sites in RNA nucleotides obtained from CLIP data as well as crosslinked amino acids in proteins obtained from RNA-interactome capture onto experimentally determined protein-RNA complex structures, as deposited in Protein Data Bank (PDB)^26^. The use of PxR3D-map to integrate CLIP and structure data is illustrated in Fig. 1a. For each protein-RNA complex, we determine the crosslinked nucleotide(s) in the RNA ligand by searching for CIMS/CITS in all instances of this sequence in the transcriptome. We then examine how the crosslinked nucleotides (s) interact with amino acids in the 3D complex structure to find structural features uniquely associated with these nucleotides as compared to noncrosslinked nucleotides. To automate this process, we search for various structural features associated with each nucleotide in the RNA ligand using programs DSSR and SNAP in the 3DNA software suite^27, 28^. In total, 15 groups of structural features are extracted to annotate each nucleotide, including RNA nucleotide conformation (e.g., base conformation and sugar puckering) and RNA-secondary structural features (e.g., single vs. double stranded region). The types of protein-RNA contacts considered include hydrogen bonds, planar amino-acid side-chain base stacking (π-stacking), and planar amino-acid base pairing (pseudo pairing)^29^. To facilitate machine learning-based classification of each nucleotide’s crosslinking status, the list of features is tabularized by aggregating the same type of contacts for each unique amino acid (e.g., 20 features that summarize the number of hydrogen bonds with each of the 20 amino acids). Additional features are also included by aggregating amino acids with similar properties (six categories: polar, positive, negative, hydrophobic, aromatic and aliphatic). This results in a total of 246 structural features associated with each nucleotide (Supplementary Table 1). Nucleotides from all protein-RNA complexes with both CLIP data and PDB structures are then compiled together, and the crosslinking status of each nucleotide is classified by its structural features using a random forest model, which also ranks features based on their importance for classification. Similarly, for protein-RNA complexes with crosslinked amino acids identified by RNA-interactome capture, the same strategy is applied to predict the crosslinking status of each amino acid based on its associated structure features (Supplementary Table 2).

**Figure 1:**
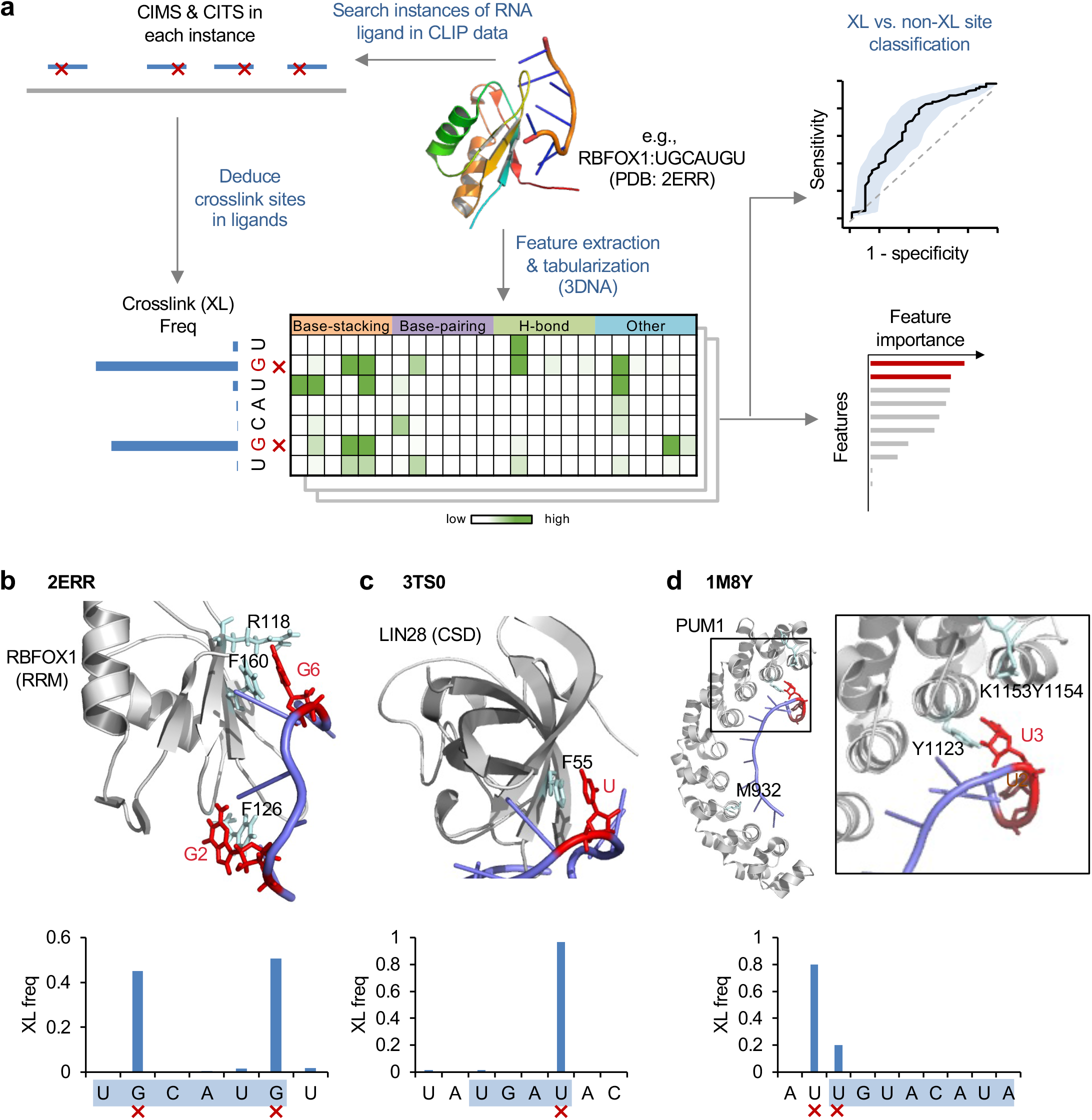
Overview of PxR3D-map to predict photo-crosslinking in native protein-RNA complexes. **a**, Schematic of PxR3D-map to predict crosslinked nucleotides in RNA using structural features. For structurally resolved protein-RNA complexes, the crosslink sites in the RNA ligand are determined by searching CIMS and CITS in instances of this sequence in CLIP data. Structural features associated with the crosslink sites including how these nucleotides contact amino acids in the complex are then examined. Specifically, the protein-RNA complex structure is analyzed by DSSR and SNAP in the 3DNA software suite to automatically extract various structural features including RNA nucleotide conformation, secondary structures and various types of nucleotide-amino acid contacts including planar amino acid sidechain-base stacking (BS), pseudo base pairing (BP), and different types of hydrogen bonds (H-bond). These structure features are tabulated based on their association with each nucleotide in the RNA ligand (e.g., how many base stacking interactions of a nucleotide with each of the 20 amino acids). These features are used to predict the crosslinking status of each nucleotide by training a random forest model, which is also used to rank feature importance for their contribution to classification. **b-d**, Three examples of protein-RNA complexes. For each complex, the bar plot shows the crosslinking frequency of each nucleotide position and the major crosslinked nucleotides are indicated at the bottom. The structure of the protein-RNA complex illustrated using PyMOL^60^ is shown at the top, with RNA in pale blue and protein in gray cartoons. The crosslinked nucleotides (red) in RNA and the crosslinked amino acids (cyan) in protein are shown in stick to highlight the nucleotide-amino acid contacts. (b) RBFOX1 RRM in complex with UGCAUGU. (c) LIN28 in complex with pre-let7-f1. (d) PUM1 in complex with AUUGUACAUA. The PDB accession code of each structure is indicated.

To test the feasibility of this strategy, we first examined several examples that 1) represent different types of RNA-binding domains (RBDs) that recognize distinct RNA sequence motifs; and 2) have crosslink sites in the RNA and protein unambiguously determined. We previously demonstrated that RBFOX, which binds the UGCAUG motif through an RNA-recognition motif (RRM), crosslinks to guanines G2 and G6 in the motif sequence by CIMS and CITS analysis^18^. Not surprisingly, G2 and G6 were identified as the predominant crosslink sites when we performed a similar search using the RNA ligand, the heptamer UGCAUGU, which was used to determine RBFOX1 RRM-RNA complex structure (PDB accession: 2ERR^30^; Fig. 1b). These two nucleotides form base-stacking with two phenylalanines, F126 and F160, respectively, which were recently identified as major crosslinked amino acids^21, 23, 31^. In addition, R118 was also found to be crosslinked in a recent study^21^, and this amino acid forms hydrogen bond with G6. In a second example, we checked the complex formed by LIN28 and precursor microRNA pre-let7-f1 (PDB accession: 3TS0; ref.^32^), with a particular focus on the region interacting with the cold shock domain (CSD). We previously determined that LIN28 CSD recognizes a UGAU motif, with crosslinking at the last uridine^33^. This is confirmed when we searched crosslink sites using the CSD binding region sequence (UAUGAUAC) of pre-let7-f1. The crosslinked uridine stacks with phenylalanine F55, which was also experimentally validated as the crosslinked amino acid^24^ (Fig. 1c). In the last example, we examined PUM1, which recognizes an 8-mer motif UGUANAUA through eight Pumilio RNA-binding repeats (Fig. 1d). When we searched CLIP data using the RNA ligand in complex structure (PDB accession: 1M8Y^34^), we found crosslinking to the first uridine (U1) of the motif and also another uridine immediately upstream (U-1). In the crystal structure, U1 stacks with a tyrosine (Y1123); U-1 does not directly contact the protein in the structure, but several amino acids in the vicinity (K1153 and Y1154) were found to be crosslinked to RNA^21^, and they may interact with U-1 in cells. These examples demonstrate a striking degree of selectivity of crosslink sites between protein and RNA, indicating requirements of certain structural features of protein-RNA contacts to induce photo-crosslinking in the cellular environment.

### Distinct structural features associated with crosslinked nucleotides

Among RBPs with both high-resolution protein-RNA complex structures deposited in PDB and in-depth CLIP data available, we were able to infer crosslink sites in the RNA ligand unambiguously for 29 non-redundant protein-RNA complexes representing 25 RBPs (Supplementary Table 3; Methods). These complexes have a total of 214 nucleotides in the RNA ligands directly contacting the proteins, including 43 nucleotides that were defined as crosslinked nucleotides and the remaining 171 as non-crosslinked nucleotides. Structural features associated with each nucleotide were extracted as described above (Supplementary Table 4).

We first performed statistical comparison of crosslinked vs. non-crosslinked nucleotides interacting with protein by examining individual sequence and structural features. We observed statistical differences in base composition between the two groups (p=0.0013, chi-squared test), with an enrichment of guanine in the crosslinked group; uridine, which is known to be susceptible to UV-crosslinking^35, 36^, is similarly enriched in the two groups (Fig. 2a). This pattern appears to be driven mainly by crosslinking to RRMs (Fig. 2b).

**Figure 2:**
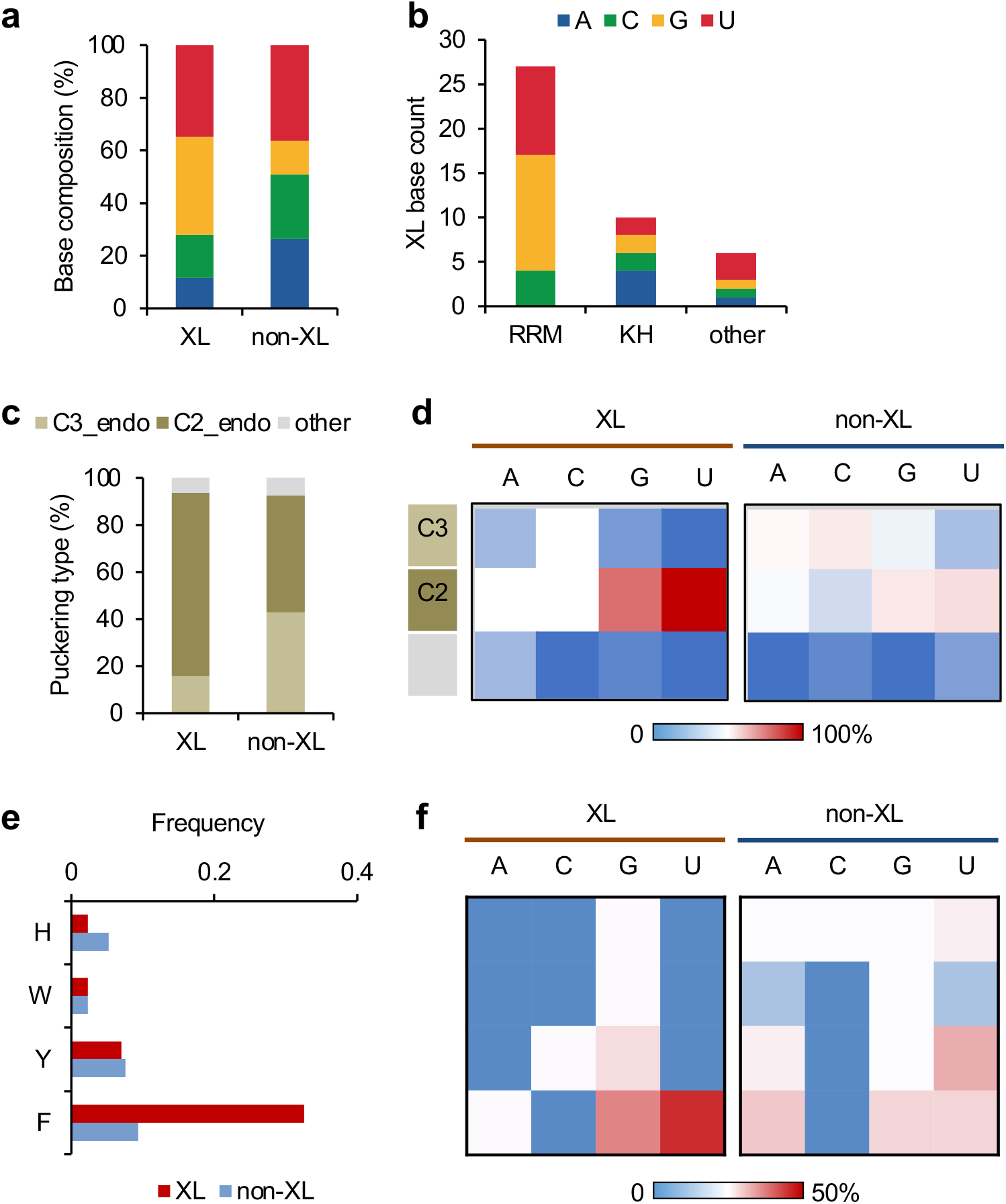
Comparison of crosslinked and non-crosslinked nucleotides using associated structural features. Only nucleotides in direct contact with amino acids were included in the analysis. **a**, Base composition of crosslinked (XL) vs. non-crosslinked (non-XL) nucleotides. **b**, Base composition of crosslinked nucleotides for RRMs, KH-domains and other types of RBDs. **c**, Distribution of sugar puckering types for crosslinked vs. non-crosslinked nucleotides. **d**, Similar to (c), but shown for each nucleotide base separately using heatmaps. **e**, Distribution of base stacking with aromatic amino acids for crosslinked vs. non-crosslinked nucleotides. **f**, Similar to (e), but shown for each nucleotide base separately using heatmaps.

RNA nucleotide conformation has been implicated to play a role in protein-RNA recognition^37^. Interestingly, the crosslinked nucleotides favor the C2’-endo conformation in their sugar puckers, while the non-crosslinked nucleotides show similar percentages in C2’-endo or C3’-endo conformation (odds ratio=4.1, p=0.005, Fisher’s exact test; Fig. 2c). In addition, the enrichment of the C2’-endo conformation is particularly prominent for crosslinked guanines and uridines (Fig. 2d). We also observed that nucleotides in direct contact with protein more frequently adopt the anti-conformation rather than the syn-conformation in the base, the preference is even more for pyrimidines (Supplementary Fig. 1a,b), which is consistent with a previous study^37^. However, there is no notable differences in base conformation between crosslinked and non-crosslinked nucleotides.

We next examined amino acids in direct contact with crosslinked vs. non-crosslinked nucleotides, but did not observe notable differences in amino acid composition between the two groups (Supplementary Fig. 1c), suggesting that the amino acid identity alone does not determine the specificity of crosslinking. In contrast, when we compared the type of protein-RNA contacts, we found a significant enrichment of base stacking with phenylalanine for crosslinked nucleotides as compared to the non-crosslinked nucleotides (0.33 vs. 0.09 events per site, p=9.5e-4, Binomial test; Fig. 2e,f). From this analysis, we did not observe notable differences in other types of amino acid-nucleotide contacts, such as hydrogen bonds (Supplementary Fig. 1d), despite their importance for determining both the specificity and affinity of protein-RNA interactions.

### Prediction of crosslinked RNA nucleotides based on structural features

To more systematically assess the contribution of different structural features and their combinations to photocrosslinking in protein-RNA complexes, we applied random forest-based classification models to predict the crosslinked vs. non-crosslinked nucleotides using structural features. For this analysis, the performance of the model was evaluated by 10-fold cross-validation. Notably, structural features are clearly predictive of crosslinked vs. non-crosslinked nucleotides with an AUC (area under ROC curve) of 0.69 (95% confidence interval between 0.60 and 0.79; Fig. 3a and Supplementary Table 4). We confirmed that this performance is robust with regard to the choice of a wide range of model parameters (Supplementary Fig. 2). In addition, including nucleotide and overlapping di-nucleotide identities further increased AUC to 0.74 (95% confidence interval between 0.65 and 0.83; Supplementary Fig. 3a), consistent with the difference of crosslinked vs. noncrosslinked nucleotides in base composition, as we identified above.

**Figure 3:**
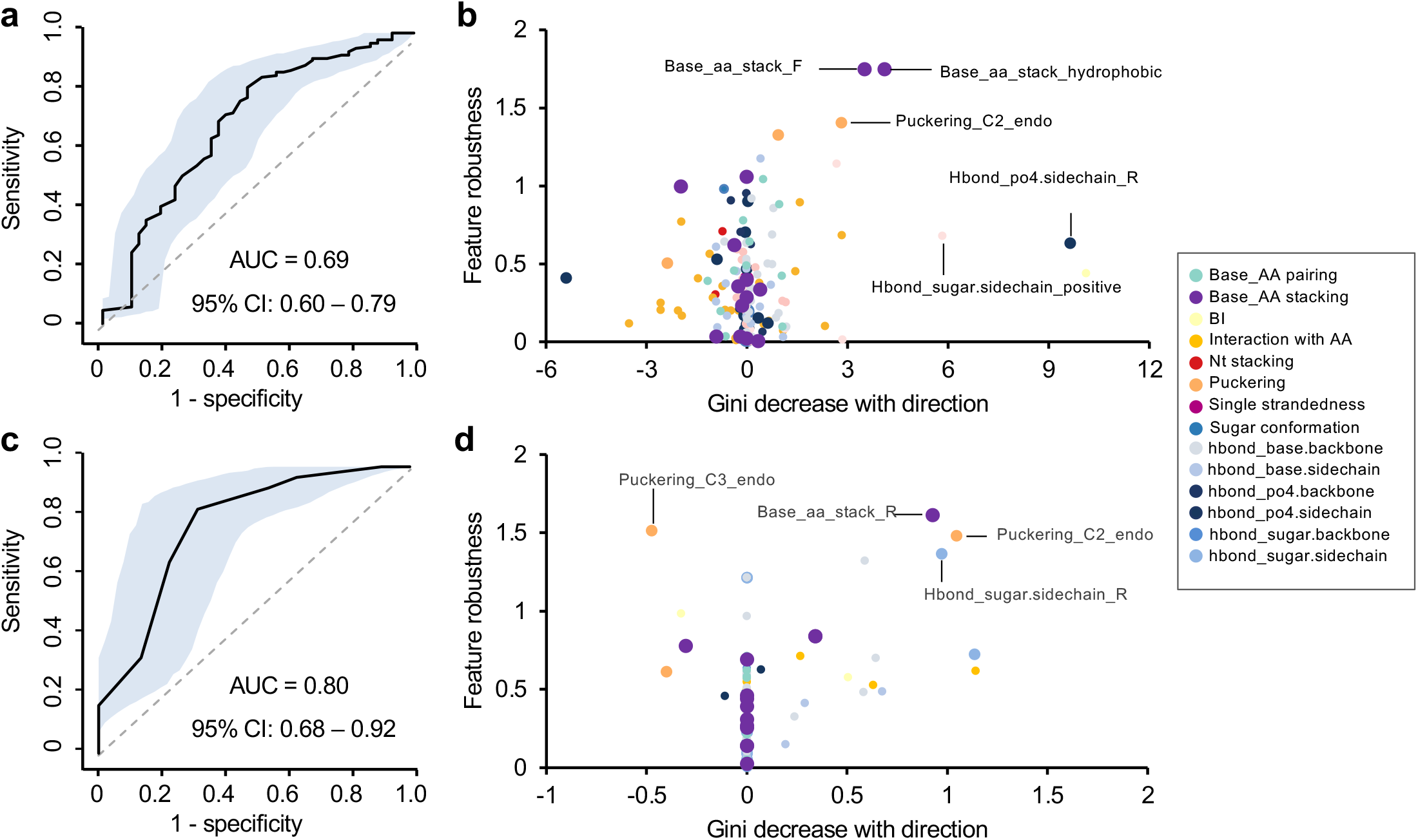
Prediction of crosslinked vs. non-crosslinked nucleotides using random forest. **a**, Prediction performance of crosslinked vs. non-crosslinked nucleotides as measured by AUC (black curve). The shaded area indicates 95% confidence interval as determined by 2000 models trained with bootstrapped data. **b**, Feature importance plot with Mean GiniDecrease of each feature shown in x-axis and feature robustness derived from permutation tests shown in y-axis. The direction of Mean GiniDecrease represents whether the feature is positively or negatively associated with crosslinking. Different feature groups are color-coded. **c, d,** Similar to (a,b) but the analysis is limited to crosslinked vs. non-crosslinked nucleotides stacking with aromatic amino acids.

Random forest models provide ranks of features based on their importance for classification reflected in reduction in Gini index^38^. Since our sample size used for classification is relatively small, we took caution to ensure the ranks of features are robust using permutation tests. In addition, we also used a generalized linear regression model to determine whether each feature contributes positively or negatively to crosslinking, which was not provided by the random forest model (see Methods). Consistent with results obtained from analysis of individual features, we found that planar base stacking with phenylalanine, or hydrophobic amino acids as a group, and the C2’-endo conformation of sugar puckering represent the top-ranked features that facilitate crosslinking (Fig. 3b, Supplementary Fig. 3b and Supplementary Table 5). Interestingly, there is also indication that certain types of hydrogen bonds, such as those formed between the phosphate (PO4) group and the sidechain of arginine, may contribute to crosslinking.

Together, our random forest prediction as well as analysis of individual features suggests that nucleotide base stacking with aromatic residues represents a prominent structural feature that facilitate crosslinking. However, we also noticed that only 22 of 76 (29%) stacking interactions had detected crosslinking, while the remaining 54 (71%) cases did not. To investigate additional structural features that may contribute to crosslinking cooperatively with base-stacking, we focused on the 76 nucleotides stacking with aromatic residues, and built another random forest predication model. In this analysis, we achieved a predication accuracy AUC of 0.8 (95% confidence interval between 0.68 and 0.92; Fig. 3c and Supplementary Fig. 3c). Evaluation of feature importance for prediction suggests that the C2’-endo sugar puckering type, as well as interaction with arginine through hydrogen bond and base stacking, contributes to protein-RNA crosslinking (Fig. 3d and Supplementary Fig. 3d). The presence of such structural features appears to be particularly common for RRM, as shown in the example of RBFOX. In this case, G6 of the UGCAUG motif interacts with phenylalanine (F160) and arginine (R118) through base stacking and hydrogen bound, respectively, and both amino acids were found to be crosslinked (Fig. 1b). Taken together, our analysis of structural features associated with crosslinked nucleotides suggests the structural requirements to induce protein-RNA photo-crosslinking and the importance of base stacking with phenylalanine, specific types of hydrogen bonds and nucleotide conformation in this process.

### Distinct structural features associated with crosslinked amino acids

To further validate our finding in the structural features associated with crosslinked nucleotides, we applied the PxR3D-map method to analyze crosslinked amino acids identified by RNA-interactome capture. Among the previous studies that reported crosslinked amino acids at single residue resolution^20–22^, we decided to focus our analysis on data obtained from RBS-ID, which represents the most comprehensive collection of crosslinked amino acids^21^. After intersecting with protein-RNA complex structures in PDB, we obtained 55 nonredundant complexes (Supplementary Table 6), which consist of 116 crosslinked amino acids and 1,380 non-crosslinked amino acids that are in direct contact with RNA. For each of these amino acids, we extracted 36 structural features describing the identity of the interacting nucleotides, and the type of protein-RNA contacts including hydrogen bonds, planar side-chain base stacking, and pseudo pairing (Supplementary Tables 2 and 7).

Among all crosslink sites identified by RBS-ID without filtering by protein-RNA complex structures, the most abundant amino acids reported were cysteine, followed by phenylalanine, tyrosine and arginine. The enrichment of phenylalanine, tyrosine, and arginine is consistent with our analysis on amino acids interacting with crosslinked nucleotides, as described above. On the other hand, cysteine was not enriched among amino acids directly contacting crosslinked nucleotides, although it was previously shown that cysteine was susceptible for UV crosslinking with nucleotides^15, 39^. To resolve this discrepancy, we examined the proportion of crosslinked amino acids in all types of annotated RBDs. Interestingly, while 67% of crosslinked phenylalanine and 43% of crosslinked tyrosine are located in annotated RBDs, the proportion is much lower for cysteine (29%) (Fig. 4a). When we focused on crosslinked amino acids located in RBDs with protein-RNA structures, the proportion of crosslinked amino acids directly contacting RNA is also much lower for cysteine (4/16=25%), as compared to phenylalanine (29/34=85%) and tyrosine (5/6=83%) (Fig. 4b). Overall, among amino acids that directly contact RNA in structurally resolved protein-RNA complexes, crosslinked amino acids are mostly enriched for aromatic residues phenylalanine, tyrosine, and tryptophan, followed by a relatively moderate enrichment for methionine and cysteine (Fig. 4c). We also examined the base composition of the closest nucleotides each crosslinked vs. non-crosslinked amino acid directly contacts, as a proxy of the nucleotide it might crosslink to. In this analysis, the base compositions for the two groups are in general similar (p=0.25, chi-squared test), with only a slight enrichment of G/U for crosslinked amino acids (Fig. 4d). Finally, we examined the types of amino acid-nucleotide contacts, and found that the only type of contacts enriched in crosslinked vs. non-crosslinked amino acids is base stacking (0.53 event vs 0.13 per amino acid; Fig. 4e). This enrichment was observed for all four aromatic residues (Fig. 4f) and they do not appear to show an overt bias for particular nucleotides, as compared to non-crosslinked amino acids interacting with RNA (Fig. 4g). This analysis confirmed that base stacking between aromatic residues and RNA nucleotides can strongly facilitate photo-crosslinking.

**Figure 4:**
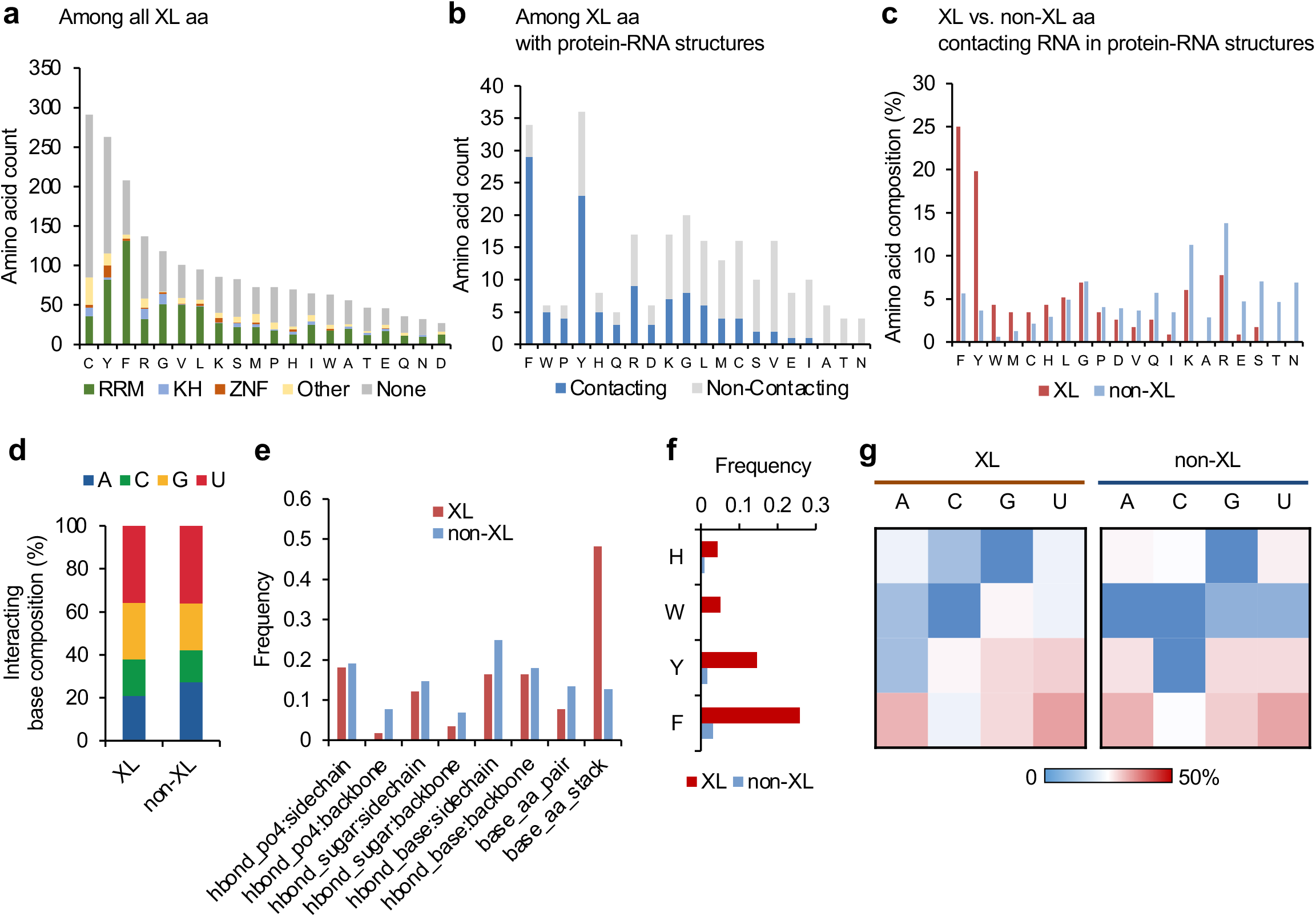
Comparison of crosslinked and non-crosslinked amino acids using associated structural features. **A,** The number of crosslink sites grouped by amino acid and RBD types. Amino acids are ordered by the total number of crosslinked sites. **b**, Among the crosslinked amino acids that can be mapped to structurally resolved protein-RNA complexes, the number of crosslink sites with and without direct RNA contacts is shown. Amino acids are ranked by the proportion of RNA-contacting crosslink sites. c, Among all RNA-contacting amino acids in structurally resolved protein-RNA complexes, the amino acid composition is shown for crosslinked and non-crosslinked sites, separately. Amino acids are ranked by enrichment in crosslinked vs. non-crosslinked sites by a binomial test. **d**, Among all RNA-contacting amino acids in structurally resolved protein-RNA complexes, the base composition of the closest nucleotides is shown for crosslinked and non-crosslinked sites separately. **e**, Frequency of structural features associated with crosslinked and non-crosslinked amino acids. **f**, Frequency of base stacking interactions for aromatic residues at crosslinked and non-crosslinked sites. **g**, Similar to (f) but shown for individual nucleotide bases separately in heatmaps.

### Prediction of crosslinked amino acids based on structural features

We next focused on the 116 crosslinked and 1,380 non-crosslinked amino acids that are in direct contact with RNA and applied PxR3D-map to predict crosslinked vs. non-crosslinked amino acids using structural features. A random forest model was trained using the amino acid and dipeptide identities as well as the 36 structural features describing amino acid-nucleotide contacts. In this analysis, we achieved an AUC of 0.76 (95% confidence interval between 0.71 and 0.82; Fig. 5a and Supplementary Table 7). Again, the performance of the random forest model is robust with respect to a wide range of model parameter choices (Supplementary Fig. 4). By applying the same feature ranking method as described above, we found aromatic residues including phenylalanine and tyrosine as well as stacking of these residues with nucleotide bases, especially uridine and guanine, as the most strongly associated features for prediction. Interestingly, we found that several dipeptides involving glycine, such as GF and GL, are also ranked among the top features (see below). Consistent with our previous analysis of crosslinked nucleotides, most types of hydrogen bonds seem to have no or only moderate contribution to facilitating crosslinking (Fig. 5b and Supplementary Table 8).

**Figure 5:**
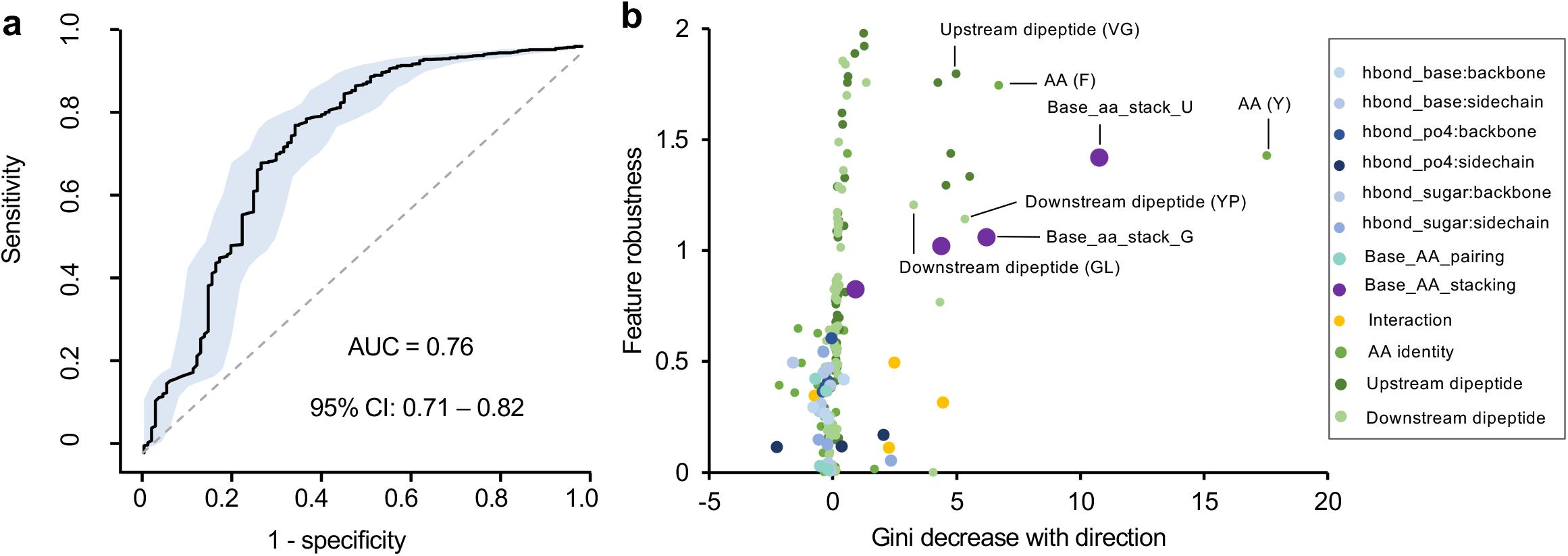
Prediction of crosslinked vs. non-crosslinked amino acids using random forest. **a**, Prediction performance of crosslinked vs. non-crosslinked amino acids as measured by AUC (black curve). The shaded area indicates the 95% confidence interval as determined by 2000 models trained with bootstrapped samples. **b,** Feature importance plot with Mean GiniDecrease of each feature shown in x-axis and feature robustness derived from permutation tests shown in y-axis. The direction of Mean GiniDecrease represents whether the feature is positively or negatively associated with crosslinking. Different feature groups are color-coded.

### Different mechanisms of photo-crosslinking for RRMs and KH domains

Taking advantage of the expanded number of crosslink sites identified by RNA-interactome captures, we next examined whether different types of RBDs have different crosslinking mechanisms. It is well known that different RBDs have distinct amino acid compositions and structural characteristics at the protein-RNA interaction interface. For example, RRMs and KH-domains are the two most abundant types of RBDs with distinct folds and RNA binding specificities. While aromatic residues, which we found to facilitate crosslinking, are frequently found in RRMs, they are rarely present in KH domains^40, 41^.

By comparing the residue composition of crosslinked amino acids in different types of RBDs, we found aromatic residues, such as phenylalanine and tyrosine, are indeed the most abundant at the crosslink sites in RRMs. This is in stark contrast to KH domains, in which crosslinking frequently occurs at cysteine, arginine, and glycine. For zinc finger (ZNF) domains, tyrosine is the most represented (Fig. 6a).

**Figure 6:**
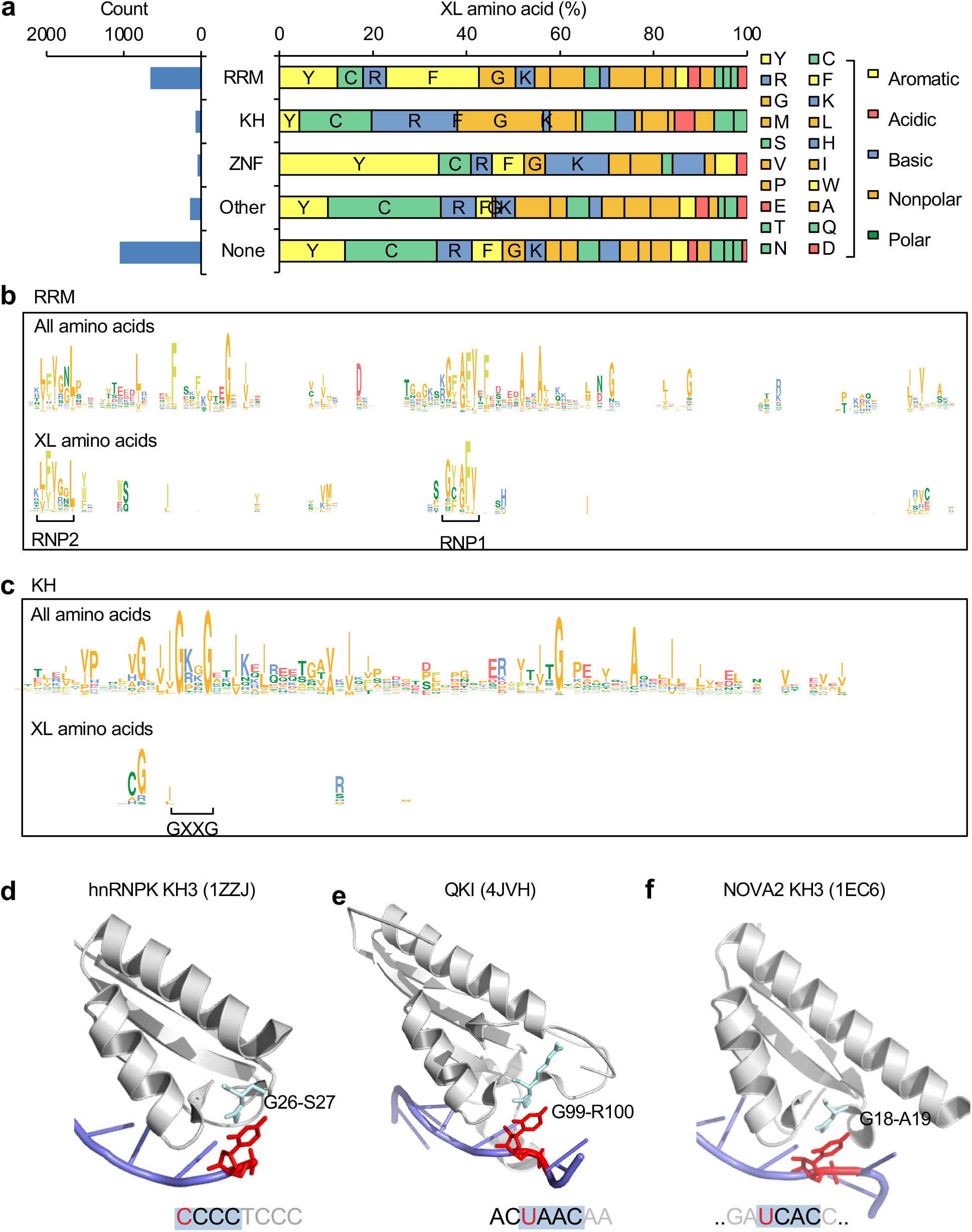
Distinct protein-RNA crosslinking mechanisms for different types of RBDs. **a**, The number (left) and composition (right) of crosslinked amino acids in different types of RBDs. **b**, Top: Multiple sequence alignment logo for RRMs. The total height of each position shows the information content of all amino acids in each position, while the relative height of each letter reflects the amino acid composition at that position. Amino acid color codes are the same as in (a). Bottom: Crosslinked amino acids at each position shown in the same format as in the top. Note crosslinked amino acids are visible only for positions with non-zero information content. The two RNP motifs defining RRMs are indicated. **c**, Similar to (b) but for KH-domains. The GXXG motif defining the KH-domains is indicated. **d-f**, Examples of KH-domains in which a glycine-related dipeptide stacks over the base of the first position of its tetramer RNA sequence motif. In each panel, The structure is illustrated using PyMOL^60^, with protein in gray and RNA in blue cartoons. The RBD and PDB accession code are indicated. (d-e) The GX dipeptide shown in cyan (stick) indicates crosslinked amino acids. The dipeptide bond stacks over the first position (red) of the tetramer RNA motif, highlighted using a shaded box in the RNA nucleotide sequence. (f) The first nucleotide (red) of the UGAC motif represents the predominant crosslink position determined using CLIP data. The GA dipeptide interacting with this uridine is highlighted in cyan (stick).

To investigate unique mechanisms of protein-RNA crosslinking that may underlie specific types of RBDs, we focused on the comparison of RRMs and KH domains. For each type, we compiled a list of domains, including those with crosslink sites identified. Multiple sequence alignments were then performed, so that the crosslink sites in different domains of each type could be compared (Supplementary Figs. 5 and 6). This analysis confirmed that the major crosslink sites for RRMs are located in the two ribonucleoprotein domains RNP-1 and RNP-2, which are two defining features of RRMs that are critical for RNA binding. There, the two conserved phenylalanines are the most frequent crosslink sites (Fig. 6b), and they typically form base stacking with RNA (Fig. 1b and additional examples discussed in ref.^20, 21, 23^). In contrast, for KH domains, the most crosslink sites are located adjacent to the GXXG (X=any amino acid) motif, a defining feature of the KH domain, in the N-terminus; these crosslink sites frequently involve a dipeptide including glycine, such as cysteine-glycine (Fig. 6c). In general, cysteine is crosslinked in the context of the cysteine-glycine dipeptide, while glycine is crosslinked in GX or XG dipeptide in the absence of cysteine. However, crosslinking rarely occurs in the GXXG motif.

KH-domain is known to recognize an RNA tetramer motif, such as YCAY (Y=C/U) for NOVA^41–44^. When we examined these protein-RNA complex structures in detail, we found that the dipeptide at the crosslink site interacts with the first position of the tetramer. For example, the glycine has evidence of crosslinking in hnRNP K KH3 and QKI STAR domains (a variant of KH domains with two flanking Qua domains)^21^, and they interact with the first cytosine of the CCCC motif for hnRNP K^44^ (Fig. 6d) and the first uridine of the UAAC motif for QKI (Fig. 6e; in this case the first two nucleotides in the QKI binding sequence motif ACUAAC are recognized by the Qua2 domain)^45^, respectively. In the case of NOVA, the crosslink site in protein is unknown. However, we previously determined that the first uridine in the YCAY motif represents the predominant crosslink site in RNA. This nucleotide contacts a glycine-alanine dipeptide in NOVA2 KH3 (G18A19, PDB accession: 1EC6; ref.^43^; Fig. 6f). Importantly, in all these cases, the GX or XG dipeptide bond appears to stack over the base of the contacting nucleotide^46^. This is confirmed by a systematic search for planar stacking formed between dipeptide bonds and nucleotide bases using 3DNA-SNAP^27, 28^. Furthermore, our survey also found additional examples including NOVA1 KH3 (G18A19; PDB accession: 2ANN^46^), PCBP2 KH1 (G26S27; PDB accession: 2PY9, ref.^47^) and KH3 (G300-S301, PDB accession: 2P2R^48^). This analysis suggests that crosslinking of KH domains with RNA utilizes a distinct mechanism from base stacking over aromatic residues observed from RRMs.

## Discussion

In this study, we developed a computational method PxR3D-map to systematically analyze structural features associated with and likely contributing to photo-crosslinking between protein and RNA in their native complexes. Given the wide applications of UV-crosslinking to study protein-RNA interactions and their functional consequences in gene regulation, the identification and characterization of the precise crosslink sites in both RNA and protein is undoubtedly a key step to better interpret data generated by these assays and understand mechanisms underlying protein-RNA complex formation.

Although current technologies do not allow simultaneous identification of crosslink sites in RNA and protein, we demonstrated that this limitation can be mitigated by the integration of crosslink sites identified separately in protein or RNA with protein-RNA complexes with experimentally resolved 3D structures. Systematic analysis of an expanding list of complexes allowed us to shed mechanistic insights into protein-RNA crosslinking. Based on our analysis, direct contacts between amino acids and nucleotides are clearly not sufficient to induce the formation of covalent bonds between the two with notable efficiency in the experimental conditions currently used to study protein-RNA complexes. Instead, this process relies on certain structural features in the complex. Even though currently resolved protein-RNA complex structures in the PDB are still limited, several structural features facilitating protein-RNA crosslinking start to emerge. Among them, the contribution of base stacking with peptide side chain, especially aromatic residues, is the most compelling. Crosslinking at such interaction sites has been experimentally validated between LIN28 CSD and its binding motif UGAU. In this case, the last uridine stacks with a phenylalanine and get crosslinked when the complex reconstituted *in vitro* was irradiated by UV^24^.

While this manuscript was in preparation, a study was published that characterized the crosslinking of RBFOX1 RRM with the cognate binding sequence UGCAUGU in detail^23^. The authors combined mass-spectrometry analysis with isotope labeling of specific nucleotides to determine crosslink sites in RNA and protein simultaneously. This study validated our previous finding from CIMS/CITS analysis that the two guanines in the motif are crosslink sites in RNA^18^, while also determining the two phenylalanine residues that stack with the two guanines as crosslink sites in the protein. By manual examination of aromatic residues and base stacking around crosslink sites determined by RNA-interactome capture, they independently reached the conclusion on the importance of stacking between the aromatic ring and the nucleotide base for photo-crosslinking. While the exact mechanisms of peptide-nucleotide adduct formation through base stacking are yet to be established, possible mechanisms have been proposed^23^. We note that it has been known for decades that UV can induce the formation of U-U or U-C dimers for nucleotide bases stacked together^49^, probably through similar mechanisms.

Our systematic and unbiased study, which is made possible by automatic analysis of protein-RNA complex structures and structural feature extraction followed by rigorous machine learning-based classification, allowed us to identify additional structural features associated with crosslink sites. Among them, the most notable finding is the crosslinking of dipeptides most likely through nucleotide base stacking over the dipeptide bond. A systematic search for such dipeptide bonds revealed that they are most frequently associated with glycine, probably because of the small size and structural flexibility of this amino acid. This mechanism appears to be particularly important for crosslinking of KH domains with RNA. In addition, we also noticed that several other residues including valine and lysine are frequently crosslinked in RNP-1 and RNP-2 of RRMs. Moreover, certain types of hydrogen bonds, as well as the C2’-endo type of nucleotide conformation, may also contribute to crosslinking. This observation is consistent with previous work which demonstrated that DNA in the C2’-endo conformation is less UV resistant compared to the C3’-endo conformation^50^. Altogether, we propose that multiple mechanisms underly the selective photo-crosslinking between protein and RNA, which will await further validation and characterization.

Knorlein *et al.* reported that up to 78% of crosslink sites in structurally resolved protein-RNA complexes can be explained by base stacking with aromatic residues^23^. This estimate appears to be rather high based on our analysis, even when we focused on RRMs, in which aromatic residues are overrepresented at the protein-RNA interaction interface (Fig. 6b). Moreover, crosslinking through base stacking with aromatic residues is clearly not the main mechanism for KH domains, which are the second most common type of RBDs (Fig. 6c). Along this line, our analysis suggests that stacking between RNA base and aromatic residues is insufficient to induce crosslinking. A striking example illustrating this point is PUM, which recognizes each of the eight nucleotides in its RNA sequence motif through base stacking, and yet crosslinking occurs specifically at the first uridine of the motif or one nucleotide further upstream (Fig. 1d). This is also in line with our finding that additional structural features, including hydrogen bonds between the sugar of the nucleotide and the side chain of arginine, as well as C2’-endo sugar puckering, can classify crosslinked vs. non-crosslinked stacking interactions (Fig. 3c, d).

Our analysis provided insights into crosslinking of cysteine, which is the most photoreactive among amino acids due to the presence of sulfur as an electron donor^15, 39^. While cysteine is the most abundant amino acid at crosslink sites identified by RNA-interactome capture^21^, a large proportion of them are not located in known RBPs. Even for those located in the RBDs of RBPs, they frequently do not directly contact RNA, as judged from available protein-RNA complex structures (Fig. 4a,b). These observations suggest crosslinking of cysteine with RNA can occur independent of stable protein-RNA interactions. Interestingly, we note that among the 13 crosslink sites found in KH domains that involve cysteine, 10 (77%) occurs in the CG or GC dipeptide context. This proportion does not seem to be explained by overrepresentation of such dipeptides in KH domains in general, and may reflect the importance of these dipeptides to initiate crosslinking at cysteine.

It is our hope that the results of this study may provide directions to better interpreting data generated by UV-crosslinking-based assays including CLIP and RNA-interactome captures. One cautionary note is crosslinking bias, which has been discussed in the literature^51^, but the extent and underlying mechanisms are not clear. Following our analyses, such bias can arise when the structural requirement to induce crosslinking cannot be fulfilled in protein-RNA contacts that determine specific protein-RNA interactions, but is rather fulfilled through additional protein-RNA contacts which might be more transient. We previously found cases in which the crosslink sites are located in the vicinity but not part of the RBP binding motifs, including crosslinking of the upstream uridine in PUM binding motif (Fig. 1d). As another example, SRSF1 binds to GGAGGA or the half site GGA, but crosslinking occurs predominantly in an upstream uridine in the sequence UGGA and the uridine does not contribute to binding specificity or affinity^52^. Therefore, interpretation of protein-RNA interactions mapped by these assays can benefit from validation using independent approaches such as assessment of binding specificity by the presence of the consensus motifs or experimental mutagenesis.

Indeed, crosslinking of transient protein-RNA interactions without stable complex formation could be widespread. This is in line with the overrepresentation of cysteine among crosslinked amino acids with a majority lacking evidence of direct protein-RNA contacts from stable complexes, as described above. In addition, we noticed that among crosslink sites in RNA identified by CIMS and CITS analysis of RBFOX CLIP data^18^, a majority (55-60%) are uridine when the consensus UGCAUG or UGCAUG-like (with one mismatch) sequences are not present. On the surface, this observation appears to be inconsistent with the predominant crosslinking of RBFOX with guanines when stable complexes are formed between RRM and the UGCAUG element. However, this apparent discrepancy can be resolved if one envisions that crosslinking could occur when the protein scans through RNA before a cognate binding site is found. In this context, multiple interactome capture studies aimed to identify the crosslinked peptide-nucleotide adducts reported that the crosslinked RNA moiety is predominantly uridine^20, 21, 25^ and Bae *et al.* decided to focus on uridine exclusively in their mass-spec analysis to limit the search space^21^. On the other hand, it is clear that crosslinking is not limited to uridine, as demonstrated in the example of RBFOX and a number of other protein-RNA complexes^52^. Our analysis implies that some of the crosslink sites with uridine adducts could reflect transient protein-RNA interactions which might be outnumber stable complexes. Therefore, mass-spec searches for peptides conjugated with non-uridine residues may lead to discoveries of additional crosslink sites in stable protein-RNA complexes.

Finally, the specific structural requirement for protein-RNA crosslinking suggests that the crosslink sites mapped by CLIP or RNA-interactome capture could provide useful constraints when one develops models of protein-RNA complex structures. This direction is particularly encouraging because of recent progress made in computational prediction of protein structures using primary sequences^53^. These models are typically trained with large datasets, a requirement difficult to meet for protein-RNA complexes (~2000 structures in total as of June 2022). It is possible that the demands on a large number of training complexes could be mitigated by providing structural constraints informative for protein-RNA complexes as suggested by selective protein-RNA crosslink sites. In addition, we expect that experimental mapping of protein-RNA interactions can benefit from technologies that are able to crosslink protein and RNA with less selective structural requirement to achieve improved sensitivity and reduced bias. The methodological framework and analyses presented in this work could provide a guide for future development of such technologies.

## Acknowledgements

We thank Judith Kribelbauer for helpful discussion during the early stages of the project. This work was supported in part by grants from the National Institutes of Health (NIH) (R01GM124486, R01GM136856 and R03HG009528 to CZ and R01GM096889 to XJL). High-performance computation was supported by NIH grants S10OD012351 and S10OD021764.

## Author contributions

Conceptualization and experimental design: H.F. and C.Z. Data analysis: H.F., X-J.L, L.L., D.U., and C.Z., Writing: H.F. and C.Z. All authors critically reviewed the manuscript.

## Methods

### Compilation of protein-RNA complex structures and structural feature extraction

Our search for RBPs with both CLIP data and protein-RNA complex structures as deposited in Protein Data Bank (PDB)^26^ was initially performed in Jan 2018. For systematic analyses of protein-RNA complex structural features described in this study, we downloaded and parsed all PDB structures to identify macromolecular complexes involving both protein and RNA chains in July 2020. In total we identified 1,090 complexes, including 277 (human), 28 (mouse), and 6 (rat) complexes, respectively.

All these complexes were analyzed by the programs DSSR and SNAP in the 3DNA software suite^27, 28^ to determine nucleotides and amino acids in direct contacts (defined by the shortest distance between a pair of heavy atoms in the nucleotide and the amino acid ≤4.5 Å). In addition, DSSR and SNAP extracted structural features that describe RNA nucleotide conformation (e.g., base morphology and sugar puckering), RNA-secondary structures (e.g., single vs. double stranded region), and various types of RNA-protein contacts including different types of hydrogen bonds, planar amino acid sidechain-base stacking, and pseudo pairing. To facilitate downstream analysis, we summarized these features with respect to their association with each individual nucleotide in the RNA ligand (e.g., the number of hydrogen bonds with each of the 20 amino acid; 246 features in total; Supplementary Table 1) or each individual residue in the protein chain (e.g., the number of hydrogen bonds with each of the four RNA bases; 36 features in total; Supplementary Table 2) at the protein-RNA interaction interface. Additional features were also included by aggregating amino acids with similar properties (6 categories: polar, positive, negative, hydrophobic, aromatic and aliphatic; Supplementary Table 4).

### Mapping crosslink sites to structurally resolved protein-RNA complexes

To map crosslinked nucleotides of RNA ligands in structurally resolved protein-RNA complexes, we intersected the RBPs in these complexes and those with CLIP data and obtained a list of 41 RBPs^18, 54–56, 57, 58^. Crosslinking sites in RNA were mapped by crosslinking-induced mutation site (CIMS) and truncation site (CITS) analysis in our previous studies (e.g., ref.^52^) or performed in this study using the CLIP Tool Kit (CTK) package^19^. For each RNA ligand, we then searched instances of the RNA ligand sequence in the respective CLIP data to determine the frequency of crosslinking at each nucleotide position. When the RNA ligand is long and only part of the sequence interacts with RBP, the sub-sequence that resembles the consensus RNA-binding sequence motif of the RBP was used for the search. In these cases, searches with one or two nucleotide extension on each side were also performed to identify additional crosslink sites immediately flanking the consensus motif sequence. A nucleotide in the RNA ligand was defined as crosslinked nucleotide if we found ?20 instances of the ligand (or its subsequence interacting with the protein) with crosslink sites determined by CLIP, and the crosslinking frequency at the position among all instances was ?0.3. Alternatively, when a smaller number of instances with crosslinking evidence were available (?10), we required the crosslinking frequency at the position ?0.5. The remaining positions of the RNA ligand interacting with the protein were defined as non-crosslinked nucleotides (Supplementary Table 4). Positions without direct contact with amino acids were excluded in our analysis. When multiple protein-RNA complex structures were available for the same RBD, only the one supported by the most crosslinking events were kept in our analysis. We were able to determine crosslink sites in RNA unambiguously for 29 non-redundant protein-RNA complexes representing 25 RBPs, from which we obtained 43 crosslinked and 171 non-crosslinked nucleotides directly contacting the protein (Supplementary Tables 3 and 4).

To map crosslinked amino acids in structurally resolved protein-RNA complexes, we used crosslinked sites identified by RBS-ID^21^. This dataset included 1,970 crosslink sites in proteins (denoted RNA-binding sites or RBSs in the original paper) in 640 protein groups. The amino acid coordinates reported for these sites were compared with the amino acid residue coordinates in the protein chain the respective PDB structures. In total, we found 104 complex structures from 46 proteins with at least one crosslinked amino acid. We removed the redundant structures and chains by keeping the one with the maximum crosslinked peptide counts and the maximum number of non-zero structural features to obtain 55 non-redundant complexes (Supplementary Table 6). After removing amino acids without directly contacting RNA nucleotides, we obtained 116 crosslinked and 1,380 non-crosslinked amino acids used in our analysis (Supplementary Table 7).

### Prediction of crosslinked nucleotides and amino acids using random forest models

We trained a random forest model to predict crosslinked vs. non-crosslinked nucleotides using structural features as described above and prioritize the structural features that contribute to protein-RNA crosslinking. Another random forest model was developed to predict crosslinked vs. non-crosslinked amino acids and prioritize associated structural features. For these tasks, the Caret package (version 6.0.80; ref.^38^) in R was used to train the models and perform predictions.

We observed improved sensitivity without impairing specificity by adopting a sampling method that corrects the imbalance of the positive and negative samples using SMOTE^59^. Model performance was evaluated by 10-fold cross validation using ROC area under curve (AUC). To evaluate the robustness of the models with respect to model parameter choices, we trained models with different parameters including the number of trees (ntree) in the forest and the number of features per tree (mtry). For prediction of crosslinked nucleotides, the optimal performance was achieved with mtry=17 and ntree=100 as measured by AUC (Fig. 3a). Similarly, for prediction of crosslinked amino acids, the optimal performance was achieved with mtry=6 and ntree=1000 (Fig. 5a).

### Evaluation of structural feature importance

For a trained random forest model, Mean GiniDecrease was used to rank the feature importance for prediction. Given the relatively moderate sample size of data available for our classification tasks, we also developed a metric to measure the robustness of feature ranks. Specifically, we randomly permutated the crosslinking labels of the nucleotides (or amino acids) and re-trained the models with the same parameters. For each permutation and re-trained model, the features were ranked. This process was repeated for *N*=2,000 times, and the robustness of each feature is defined as 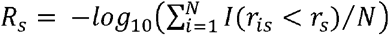 where *r_s_* is the rank of feature importance for feature *s* in the true model without crosslinking label permutation and *r_is_* is the rank of feature *s* in the *i*^th^ permutation; *I*(.) is the indictor function.

Gini index does not indicate whether a feature contributes positively or negatively to prediction of the crosslinked nucleotides or amino acids. To determine whether each feature associates with crosslinking positively or negatively, we used a generalized linear regression model using R package Caret (version 6.0.80). 10-fold cross validation was used to determine the best parameters (α=0.56 and λ=0.14 for prediction of crosslinked nucleotides; α=0 and λ=7.37 for prediction of crosslinked amino acids) that maximized the AUC. The coefficient of each feature was used to determine the contribution direction towards the prediction of the crosslinking status.

## Supplementary Figure Legends

**Supplementary Fig. 1: Characterization of crosslinked nucleotides and the associated structural features.**

**a**, Distribution of sugar conformation types for crosslinked and non-crosslinked nucleotides contacting amino acids in protein-RNA complex structures.

**b**, Similar to (a), but distributions for individual nucleotide bases are shown separately using heatmaps.

**c**, Distribution of amino acids contacting with crosslinked and non-crosslinked nucleotides.

**d**, Distribution of hydrogen bond types formed between amino acids and crosslinked vs. non-crosslinked nucleotides.

**Supplementary Fig. 2: Prediction performance of crosslinked vs. non-crosslinked nucleotides using random forest models trained with different model parameters.** A wide range of model parameters including the number of trees (ntree) in the forest and the number of features per tree (mtry) are tested, and the AUC of each model is plotted.

**Supplementary Fig. 3: Prediction of crosslinked vs. non-crosslinked nucleotides using random forest trained with nucleotide and overlapping di-nucleotide identities together with structural features.**

**A**, Prediction performance of crosslinked vs. non-crosslinked nucleotides as measured by AUC (black curve). The shaded area indicates 95% confidence interval as determined by 2000 models trained with bootstrapped data.

**B**, Feature importance plot with Mean GiniDecrease of each feature shown in x-axis and feature robustness derived from permutation tests shown in y-axis. The direction of Mean GiniDecrease represents whether the feature is positively or negatively associated with crosslinking. Different feature groups are color-coded.

**C, d,** Similar to (a,b) but the analysis is limited to crosslinked vs. non-crosslinked nucleotides stacking with aromatic amino acids.

**Supplementary Fig. 4: Prediction performance of crosslinked vs. non-crosslinked amino acids using random forest models trained with different model parameters.** A wide range of model parameters including the number of trees (ntree) in the forest and the number of features per tree (mtry) are tested, and the AUC of each model is plotted.

**Supplementary Fig. 5. Crosslinked amino acids in RRMs.**

Multiple sequence alignments of RRMs are shown with crosslinked amino acids (data from Bae et al) highlighted in red. Underscored amino acids have ≥10 spectrum counts.

**Supplementary Fig. 6: Crosslinked amino acids in KH domains.**

Multiple sequence alignments of RRMs are shown with crosslinked amino acids (data from Bae et al) highlighted in red. Underscored amino acids have ≥10 spectrum counts.

## Notes

### Competing Interest Statement

The authors have declared no competing interest.

